# Vinculin anchors contractile actin to the cardiomyocyte adherens junction

**DOI:** 10.1101/613455

**Authors:** Chelsea D. Merkel, Yang Li, Qanber Raza, Donna B. Stolz, Adam V. Kwiatkowski

## Abstract

The adherens junction (AJ) couples the actin cytoskeletons of neighboring cells to allow mechanical integration and tissue organization. The physiological demands of intercellular adhesion require that the AJ be responsive to dynamic changes in force while maintaining mechanical load. These demands are tested in the heart, where cardiomyocyte AJs must withstand repeated cycles of actomyosin-mediated contractile force. Here we show that force-responsive cardiomyocyte AJs recruit actin-binding ligands to selectively couple actin networks and promote contact maturation. We employed a panel of N-cadherin-αE-catenin fusion proteins to rebuild AJs with specific actin linkages in N-cadherin-null cardiomyocytes. In this system, vinculin recruitment was required to rescue myofibril integration and desmosome assembly at nascent contacts. In contrast, loss of vinculin disrupted junction morphology and blocked myofibril integration. Our results identify vinculin as a critical link to contractile actomyosin and offer insight to how actin integration at the AJ is regulated to provide mechanical stability and cellular organization.

## Introduction

Adherens junctions link the actin cytoskeletons of adjacent cells to provide the foundation for multicellular tissue organization. The dynamic demands of cell-cell adhesion require that the AJ be both responsive and resilient to mechanical force. This is especially true in the heart, where the AJ must transmit the mechanical forces of actomyosin contraction while maintaining adhesive homeostasis. How the AJ balances mechanical integration with contractile force to maintain tissue integrity is not clear.

Cardiomyocytes are linked through a specialized cell-cell contact called the intercalated disc (ICD). The ICD is the site of mechanical and electrical continuity between individual cardiomyocytes that allow the heart to function as a syncytium (Vite and Radice, 2014; Ehler, 2016; Vermij *et al.*, 2017). Three junctional complexes form the ICD: the adherens junction (AJ), desmosome and gap junction. The AJ and desmosome are responsible for mechanical integration by coupling the actin and intermediate filament cytoskeletons, respectively, of neighboring cells. Gap junctions permit electrical continuity through the free flow of ions. Importantly, the ICD AJ is the site of myofibril integration between cardiomyocytes and allows contractile force to be transduced across heart tissue (Luo and Radice, 2003).

The core of the AJ is the cadherin-catenin complex (Ratheesh and Yap, 2012; Mege and Ishiyama, 2017). N-cadherin, the sole classical cadherin expressed in cardiomyocytes (Kostetskii *et al.*, 2005), is a single-pass transmembrane protein with an extracellular domain that mediates homotypic, calcium-dependent interactions (Shapiro and Weis, 2009; Harrison *et al.*, 2011). The adhesive properties of classical cadherins are driven by the recruitment of cytosolic catenin proteins to the cadherin tail: p120-catenin binds to the juxtamembrane domain and β-catenin binds to the distal part of the tail. β-Catenin, in turn, recruits α-catenin to the cadherin-catenin complex. α-Catenin is an actin-binding protein and the primary link between the AJ and the actin cytoskeleton (Rimm *et al.*, 1995; Drees *et al.*, 2005; Yamada *et al.*, 2005; Buckley *et al.*, 2014; Pokutta *et al.*, 2014; Ishiyama *et al.*, 2018).

AJ binding capabilities are modified by the forces of actomyosin contraction, largely through changes in α-catenin conformation (Hoffman and Yap, 2015; Charras and Yap, 2018). Force induces a conformational change in the central M-domain of αE-catenin to reveal binding sites for ligands, many of which bind F-actin (le Duc *et al.*, 2010; Yonemura *et al.*, 2010; Choi *et al.*, 2012; Yao *et al.*, 2014; Kim *et al.*, 2015; Matsuzawa *et al.*, 2018). The force required to unfurl αE-catenin (5pN) is well within the range of a myosin motor, demonstrating the physiological relevance for this model of regulation (Finer *et al.*, 1994; Charras and Yap, 2018). The recruitment of actin-binding proteins in response to force is thought to help anchor actin to the AJ (Yap *et al.*, 2018).

Cardiomyocytes have at least two distinct actin networks at cell-cell contacts – myofibrils and the cortical cytoskeleton (Li *et al.*, 2019) – that must be integrated at AJs. Many actin-binding ligands interact with αE-catenin, including vinculin, afadin, ZO-1 and Eplin (Yonemura, 2017). In epithelia, vinculin is recruited to αE-catenin in a force-dependent manner and this interaction is thought to be important for reinforcing the αE-catenin:actin interaction (le Duc *et al.*, 2010; Yonemura *et al.*, 2010; Kale *et al.*, 2018). Likewise, epithelial afadin can also bind αE-catenin in a force-dependent manner (Matsuzawa *et al.*, 2018), where it functions to strengthen the AJ under tension (Choi *et al.*, 2016). Both vinculin and afadin localize to the ICD (Geiger *et al.*, 1985; Borrmann *et al.*, 2006) and are recruited to cardiomyocyte AJs (Li *et al.*, 2019). Vinculin is required for proper heart development and functions in cardiomyocyte adhesion and contraction (Shiraishi *et al.*, 1997; Xu *et al.*, 1998). Afadin was recently identified as having a cardioprotective role at the ICD, as mice lacking afadin were shown to be more susceptible to stress-induced injury and myopathy (Zankov *et al.*, 2017). How vinculin and afadin function in mechanical coupling at cardiomyocyte AJs is not well understood.

Here we sought to define the individual functions of αE-catenin, vinculin and afadin in coupling actin to cardiomyocyte AJs. We demonstrate that cultured neonatal cardiomyocytes recruit vinculin and afadin to AJs in a force-dependent manner, similar to epithelia. We show that loss of N-cadherin in cardiomyocytes disrupts cell-cell adhesion and dissolves junctional complexes. These phenotypes defined in our *in situ* loss-of-N-cadherin system are strikingly similar to those shown for *in vivo* models (Kostetskii *et al.*, 2005). We developed a series of N-cadherin:αE-catenin fusions to test how AJ ligand recruitment and actin-binding influences junctional complex assembly, cell contact architecture and myofibril coupling. We show for the first time that vinculin recruitment to the AJ is necessary to couple myofibrils to the developing cell-cell contacts in cardiomyocytes. Our results offer new insight into actin linkage to the AJ and identify vinculin as the key link between contractile actin networks and the cardiomyocyte AJ.

## Results

### Force regulates α-catenin ligand recruitment to cardiomyocyte AJs

Vinculin and afadin are recruited to epithelial AJs in a force-dependent manner (le Duc *et al.*, 2010; Yonemura *et al.*, 2010; Kale *et al.*, 2018; Matsuzawa *et al.*, 2018). Vinculin and afadin localize to the ICD in adult heart and proximity proteomics revealed that both are enriched at the AJ in cultured neonatal cardiomyocytes (Li *et al.*, 2019). We sought to determine if vinculin and afadin recruitment to cardiomyocyte AJs is tension-dependent. Cultured cardiomyocytes were treated with 100 μM blebbistatin to suppress myosin activity for up to one hour and stained for vinculin or afadin (Figure 1, A-D). Cardiomyocytes ceased contraction within 30 seconds of blebbistatin addition whereas DMSO did not affect contraction (unpublished observation). In blebbistatin-treated cells, both vinculin and afadin were significantly reduced at cell-cell contacts after one hour, with significant loss of vinculin seen after 30 minutes (Figure 1, E and F). This is consistent with a requirement for tension at the AJ to recruit vinculin or afadin and indicates that nascent cardiomyocyte AJs retain the ability to respond to changes in mechanical force.

**Figure 1.**
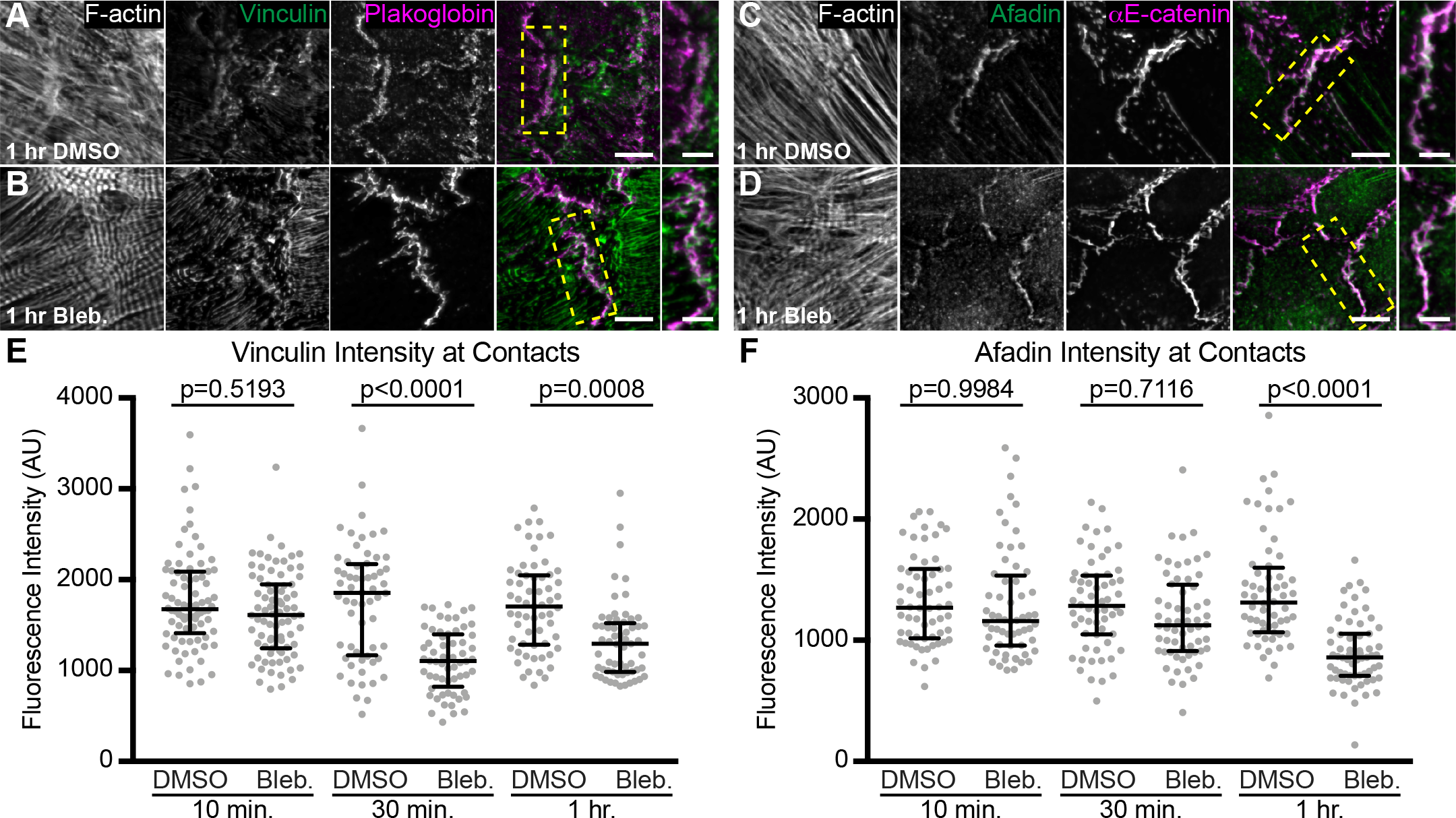
Vinculin and afadin recruitment to cardiomyocyte AJs is force dependent. A-D. Mouse neonatal cardiomyocytes were treated for 1 hr with DMSO (A or C) or 100 μM blebbistatin (B and D) before fixation. Cells were stained for F-actin (A-D), vinculin and plakoglobin (A and B), or afadin and αE-catenin (C and D). Images are max projections of 2-3 μm deconvolved stacks. Individual and merged vinculin (green) and plakoglobin (magenta) channels shown in A and B. Individual merged afadin (green) and αE-catenin (magenta) channels shown in C and D. Far right column is a magnification of the boxed contact in merge. E-F. Quantification of vinculin (E) or afadin (F) intensity at cell-cell contacts. Vinculin or afadin signal intensity was measured in cells treated with DMSO or blebbistatin for 10, 30 and 60 minutes before fixation. All data points are plotted. Middle horizontal bar is the median and error bars represent the quartile range. One-way ANOVA, n ≥ 60 images from at least 3 independent experiments. Scale bar is 10 μm in full images, 5 μm in zoomed images.

### Loss of N-cadherin disrupts cardiomyocyte cell-cell contacts

The force-responsive nature of cardiomyocyte AJs led us to question the roles of αE-catenin, vinculin and afadin in linking the AJ to actin. In order to individually test these roles, we developed a system to selectively recruit actin-binding ligands and thus control the actin-binding interfaces at the cardiomyocyte AJ. We first needed to establish a cadherin-null system in which to rebuild AJs. In intact mouse heart tissue, conditional ablation of N-cadherin causes dissolution of all AJ components as well as loss of all desmosomal and gap junction proteins at the ICD (Kostetskii *et al.*, 2005). We questioned if loss of N-cadherin would disrupt ligand recruitment and junction organization in cultured neonatal cardiomyocytes. Cardiomyocytes from N-cadherin conditional knockout mice (Ncad^fx/fx^; (Kostetskii *et al.*, 2005)) were isolated and infected with adenovirus expressing Cre-recombinase (hereby referred to as Cre).

In order to determine the time required for N-cadherin depletion post Cre-mediated recombination, we fixed Cre-infected cells at four different time points: 24, 48, 72 and 96 hours post-infection and stained for N-cadherin (Figure 2, B-F). At 24 hours post-infection, N-cadherin levels appeared similar to uninfected cells (Figure 2, B and C). At 48 hours post-infection, N-cadherin levels remained high; however, cell-cell contacts began to appear jagged with N-cadherin clustering along more linear contacts (Figure 2D). We speculate that declining N-cadherin levels are promoting AJ consolidation and altering junction morphology. Notably, at 72 and 96 hours post-infection, we observed a near complete loss of N-cadherin at cell-cell contacts (Figure 2, E and F; Supplemental Figure S1, A and B).

**Figure 2.**
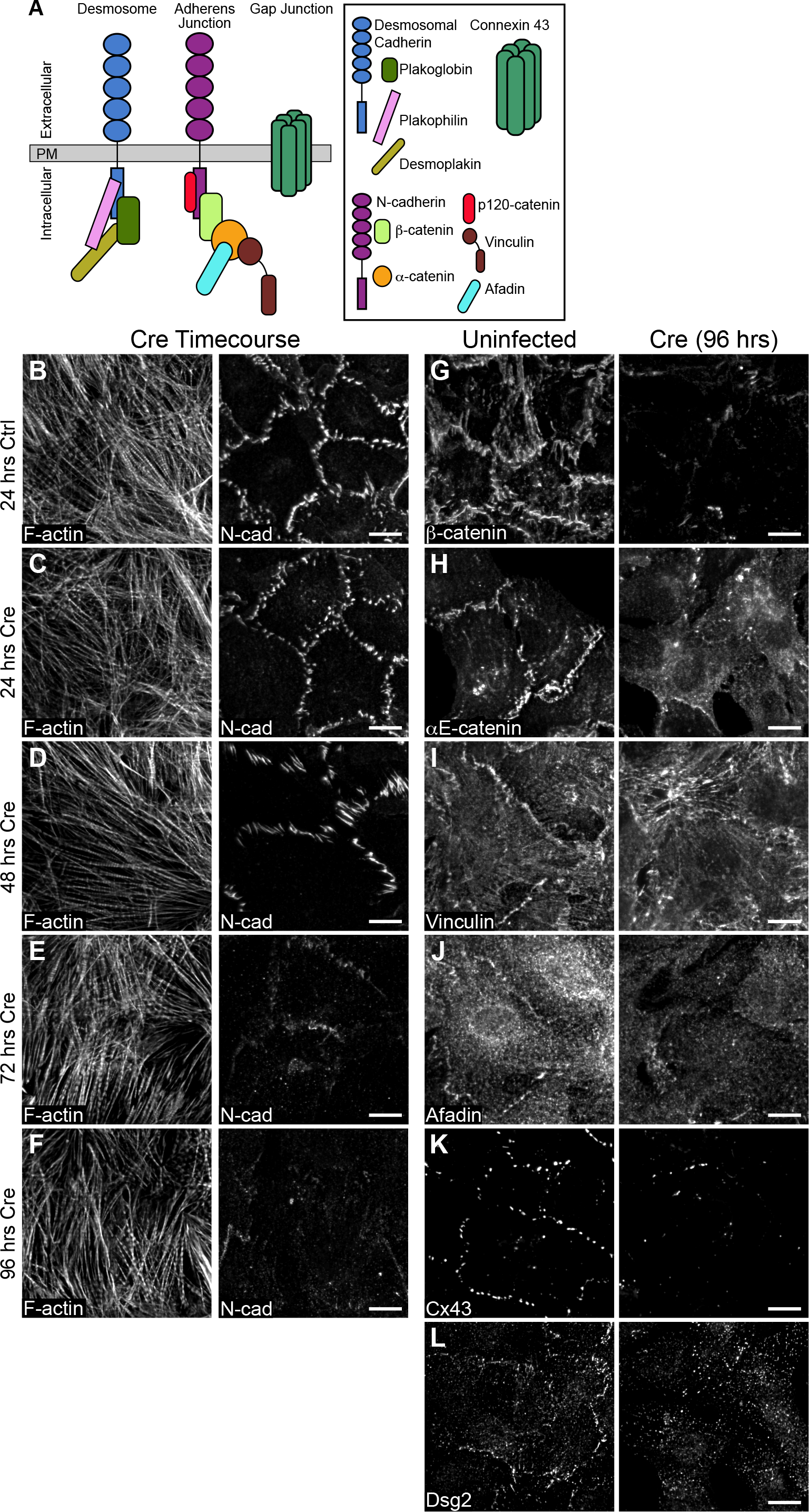
Loss of N-cadherin disrupts adhesion protein localization. A. Cartoon schematic of desmosome, AJ and gap junction proteins at cardiomyocyte cell-cell contacts. B-F. Neonatal cardiomyocytes from Ncad^fx/fx^ mice were uninfected (B) or infected with adenovirus expressing Cre recombinase (C-F) and fixed over 4 days to assess N-cadherin expression. Cells were stained for F-actin (left panel) and N-cadherin (right panel). G-L. Control and Cre-infected neonatal cardiomyocytes from Ncad^fx/fx^ mice were fixed 96 hours post-infection and stained for AJ components (G, H), AJ adapter proteins (I, J), gap junctions (K), and desmosomes (L). Images are max projections of 2-3 μm deconvolved stacks. Scale bar is 10μm.

We next assessed cell contact formation and protein recruitment at 96 hours post Cre infection. In uninfected control cardiomyocytes, AJ markers N-cadherin, β-catenin and αE-catenin were recruited to cell-cell contacts (Figure 2, G and H). Likewise, αE-catenin ligands vinculin and afadin; the gap junction protein connexin 43 (Cx43); and desmosomal markers desmoglein 2 (Dsg2, Figure 2, I-L), plakoglobin, and plakophilin 2 (Supplemental Figure S1, C and D) all localized to cell-cell contacts. In contrast, Cre expression dissolved cell-cell contacts and resulted in a loss of all AJ proteins, αE-catenin ligands, gap junctions and desmosomes (Figure 2, G-L; Supplemental Figure S1, C and D). As expected, N-cadherin is required for cell-cell adhesion in cultured cardiomyocytes and AJ formation is critical for the recruitment and organization of other junctional components.

We sought to determine if cardiomyocyte cell-cell contacts can be restored with exogenous N-cadherin-GFP. Cardiomyocytes were sequentially infected with Cre and then N-cadherin-GFP adenovirus. Expression of N-cadherin-GFP restored cell-cell contacts and the localization of AJ, gap junction and desmosome proteins (Figure 3, A-F; Supplemental Figure S1, E and F). The ability of N-cadherin-GFP to restore cell-cell contacts in an N-cadherin null background demonstrated the dynamic nature of this adhesion system and its tractability for probing cadherin and catenin function further.

**Figure 3.**
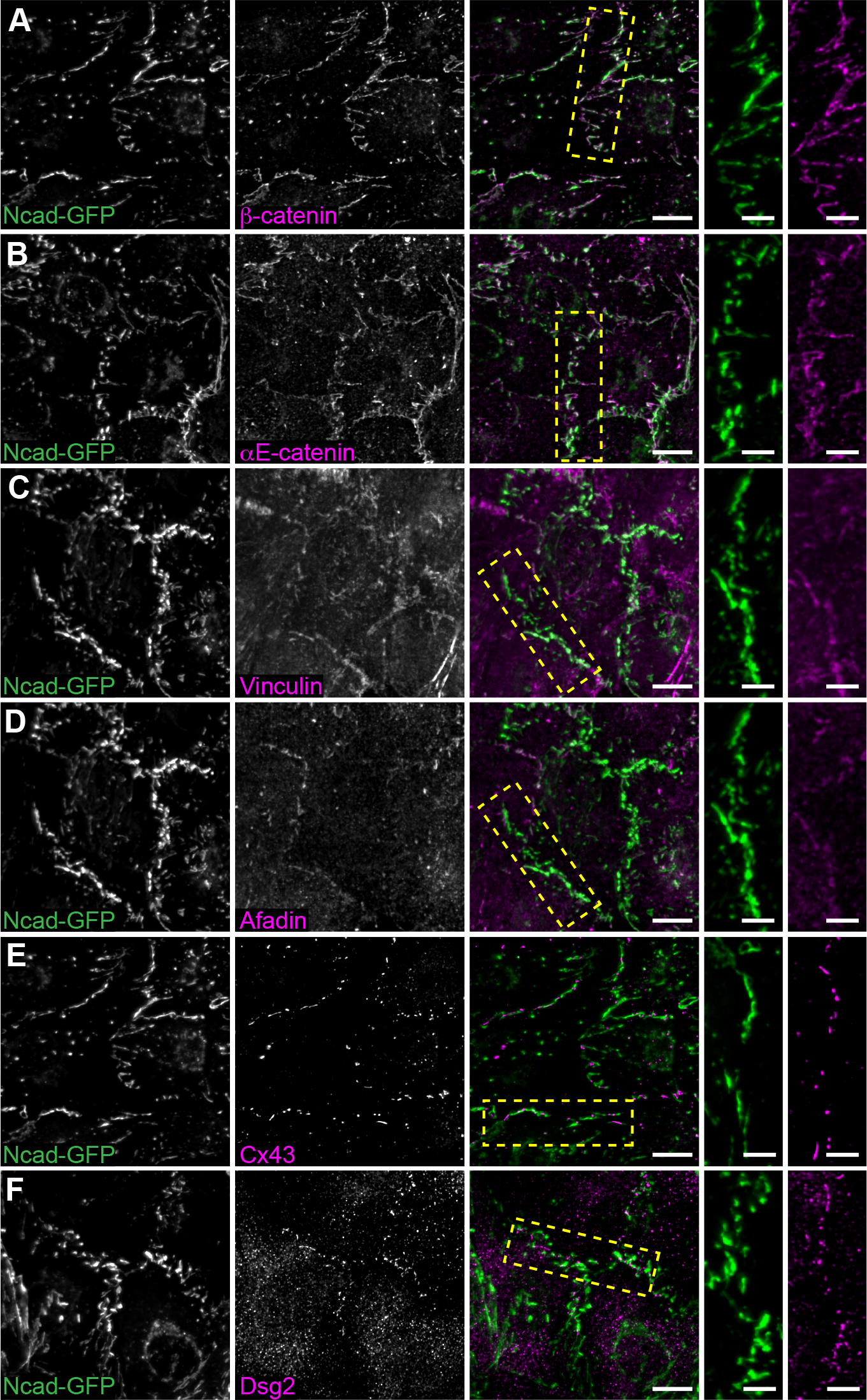
N-cadherin-GFP rescues cardiomyocyte junctional complexes. A-F. Neonatal Ncad^fx/fx^ cardiomyocytes infected sequentially with adenoviruses expressing Cre and N-cadherin-GFP, fixed and stained for AJ-associated proteins (A-D), gap junctions (E) and desmosomes (F). Individual and merged N-cadherin-GFP (green) and ICD components (magenta) channels are shown. Far right column is a magnification of the boxed contact in the merge. Images are max projections of 2-3 μm deconvolved stacks. Scale bar is 10μm in full images, 5 μm in zoomed images.

### N-cadherin-αE-catenin fusions selectively recruit αE-catenin ligands

We designed a series of N-cadherin-αE-catenin fusion constructs to systematically delineate the individual and combined functions of αE-catenin, vinculin and afadin in AJ-mediated cell-cell adhesion. Fusion constructs were created by taking the extracellular, transmembrane and p-120 binding domains of N-cadherin and fusing them to EGFP followed by the middle (M)-region and actin binding domain (ABD) of αE-catenin (Figure 4A). The M-region of αE-catenin contains three separate domains: M1, M2 and M3. The vinculin binding site is found in M1 (Choi *et al.*, 2012) and the afadin binding site spans M2-M3 (Pokutta *et al.*, 2002). The C-terminal tail of N-cadherin and the N-terminus of αE-catenin were removed to eliminate endogenous β-catenin and αE-catenin recruitment while allowing for proper N-cadherin trafficking (Davis *et al.*, 2003; Wahl *et al.*, 2003; Bianchini *et al.*, 2015). Within the N-cadherin-GFP-αE-catenin (Ncad-GFP-αEcat) fusion, we introduced various mutations or domain deletions to restrict ligand recruitment and actin-binding interfaces. Ncad-GFP-M1-ABD mimics the core cadherin-catenin complex as it possesses the αE-catenin ABD and contains both vinculin and afadin binding sites. Ncad-GFP-M1-M3 possesses the full M-region of αE-catenin but lacks the ABD and the ability to respond to tension. Ncad-GFP-M1-M2 has an open M-domain that can bind vinculin constitutively but lacks the αE-catenin ABD and afadin binding domain (Choi *et al.*, 2012). Two constructs were designed to selectively block vinculin recruitment while retaining afadin binding and actin binding through the αE-catenin ABD: Ncad-GFP-M1mutV-ABD, which contains 5 point mutations in M1 that ablate vinculin binding (Chen *et al.*, 2015), and Ncad-GFP-M2-ABD, which lacks the entire M1 domain. Note that additional ligands may bind αE-catenin M2-M3: we focus on afadin recruitment and use it as proxy for ligand binding to M2-M3. Lastly, Ncad-GFP-ABD lacks M1-M3, but retains the αE-catenin ABD.

**Figure 4.**
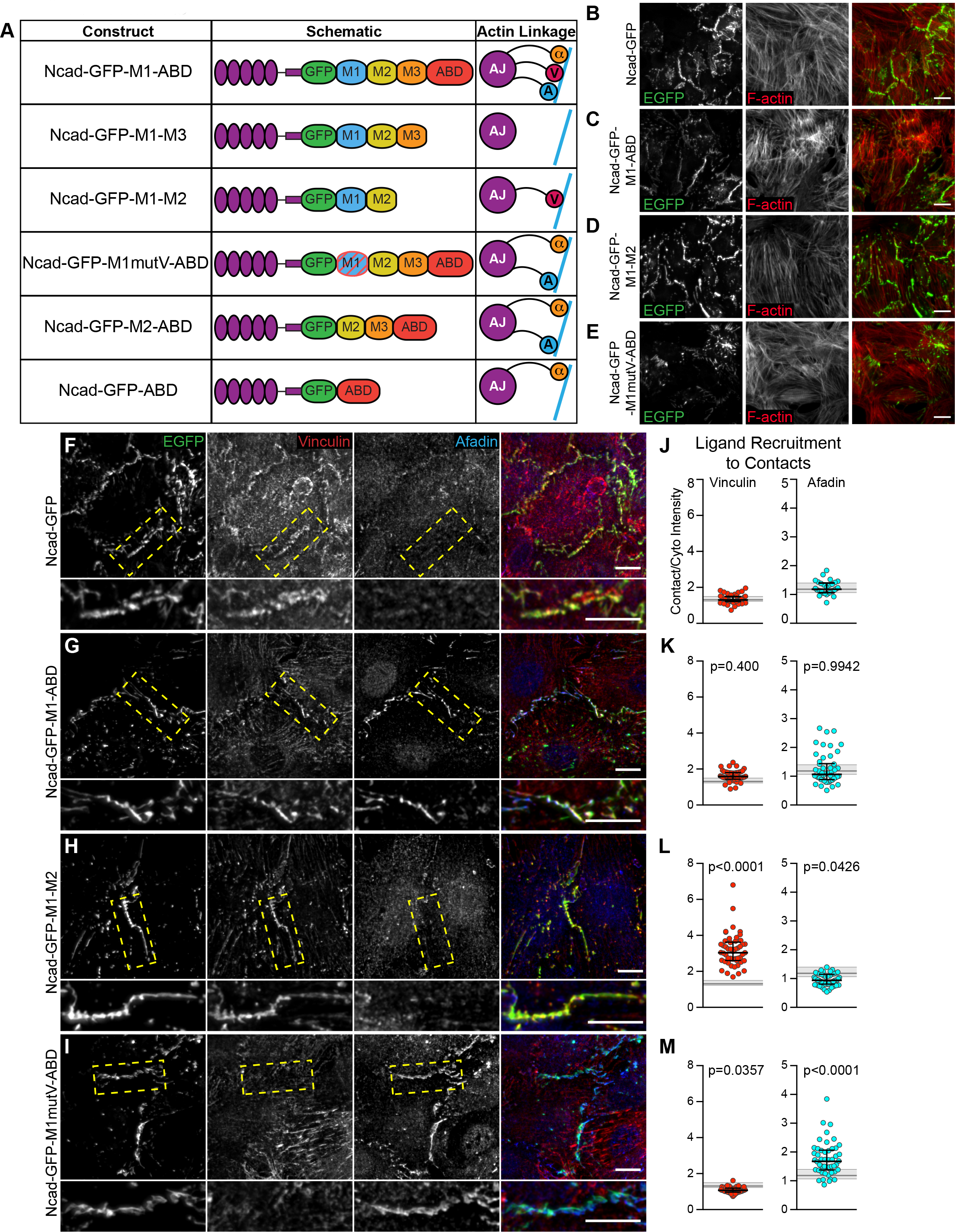
N-cadherin-GFP-αE-catenin fusions selectively recruit ligands to cardiomyocyte cell-cell contacts. A. Table of N-cadherin-GFP-αE-catenin fusion constructs used in this study. Nomenclature, domain schematic, and actin linkage cartoon shown. B-E. Lower magnification images (40X) of Ncad^fx/fx^ cardiomyocytes infected with Cre and N-cadherin-GFP (B) or N-cadherin-GFP-αE-catenin fusion adenoviruses (C-E). Individual and merged EGFP (green) and actin (red) channels are shown. Images max projections of 5 μm stacks. F-I. Cardiomyocytes infected with Cre and N-cadherin-GFP (F) or fusion adenoviruses (G-I), fixed and stained for vinculin and afadin. Individual and merged EGFP (green), vinculin (red) and afadin (blue) channels shown. Images are a max projection of 2-3 μm deconvolved stacks. Bottom image is a magnification of boxed contact. J-M. Quantification of vinculin and afadin intensities at cell-cell contacts. Signal intensity at contacts was divided by the average cytoplasmic intensity and a scatter plot of all data points is shown. The black horizontal line is the median and the error bars define the interquartile range. The shaded gray region in each plot defines the median (thick gray line) and interquartile range (thin gray lines) of vinculin or afadin recruitment observed with full-length N-cadherin-GFP (J) for comparison. One-way ANOVA, significance compared to recruitment with N-cadherin-GFP. n ≥ 50 images from at least 2 independent experiments. Scale bar is 20 μM in B-E, 10 μm in F-I and 5 μm in zoomed images in F-I

To validate fusion construct localization and ligand recruitment, we first transfected Ncad-GFP-αE-catenin fusion plasmids in to cadherin-deficient epithelial A431D cells (Supplemental Figure S2). A431D cells do not express classical cadherins but do express other components of the AJ as well as the AJ ligands vinculin and afadin. Thus, we could test the ability of the fusion constructs to restore cell-cell adhesion and selectively recruit ligands. All fusion constructs localized to cell-cell contacts and recruited the predicted ligands (Supplemental Figure S2, A-G).

We then tested the ability of Ncad-GFP-αEcat fusions to restore cell-cell contacts and selectively recruit vinculin or afadin in N-cadherin-null cells. Ncad^fx/fx^ cardiomyocytes were sequentially infected with Cre plus individual adenoviral Ncad-GFP-αEcat fusions. We observed expression and proper localization of the fusion constructs by 24 hours post-infection, which continued through 72 hours post-infection, corresponding with the maximum loss of N-cadherin (Supplemental Figure S1, G-M). All Ncad-GFP-αEcat fusions localized to the membrane and reestablished cell-cell contacts, though the gross morphology of these junctions differed markedly between constructs (Figure 4, B-E). Ncad-GFP-M1-ABD recruited both vinculin and afadin (Figure 4, G and K). This construct also allowed for formation of cell-cell contacts similar to Ncad-GFP (Figure 4, B, C, F and G), indicating that the static Ncad-GFP-αEcat fusion can substitute for the cadherin-catenin-complex. In contrast, Ncad-GFP-M1-M3, which lacked the ABD and the ability to bind actin or respond to tension, failed to recruit vinculin or afadin and formed poorly organized contacts as expected (Supplemental Figure S2H). However, the constitutively active Ncad-GFP-M1-M2 enriched vinculin, but not afadin, and restored robust cell-cell contacts (Figure 4, H and L). Ncad-GFP-M1mutV-ABD and Ncad-GFP-M2-ABD both recruited afadin, but not vinculin, and generated long, linear contacts that lacked the jagged contact morphology found in controls (Figure 4, E, I and M; Supplemental Figure S2 I). Lastly, Ncad-GFP-ABD formed poor contacts that failed to recruit vinculin but did display limited afadin recruitment (Supplemental Figure S2J). We speculate that this weak afadin recruitment to cardiomyocyte cell-cell contacts may be mediated by nectins (Satomi-Kobayashi *et al.*, 2009; Li *et al.*, 2019). Thus, we were able to specifically recruit αE-catenin ligands to nascent cardiomyocyte contacts and observed morphological changes as a function of the selective association of AJ components. Notably, Ncad-GFP-M1-M2, which only recruits vinculin, formed cell-cell contacts similar to controls whereas Ncad-GFP-M1mutV-ABD and Ncad-GFP-M2-ABD, which do not recruit vinculin, organized linear contacts (Figure 4, B and D; Supplemental Figure S2I).

To assess the effects of ligand recruitment on fusion stability along cell-cell contacts, we performed FRAP (fluorescence recovery after photobleaching) analysis of Ncad-GFP and three key Ncad-GFP-αEcat fusion constructs (Figure 5). Fluorescence recovery over 15 minutes was quantified, plotted and fit to a double exponential curve. The Ncad-GFP recovery prolife was similar to previously published data from our group (Figure 5, A and E; Li, 2019), consistent with the ability of Ncad-GFP to restore AJs in an N-cadherin-null background. The mobile fraction of Ncad-GFP and all fusion constructs was ~30%, similar to those observed for AJ components in cardiomyocytes and epithelial cells (Figure 5I; Li, 2019, Yamada, 2005). While F-actin binding is critical for AJ formation and extracellular and intracellular cadherin interactions cooperate to regulate AJ assembly (Hong *et al.*, 2013), our results suggest that AJ plaque (immobile fraction) stability is regulated by cadherin extracellular interactions.

**Figure 5.**
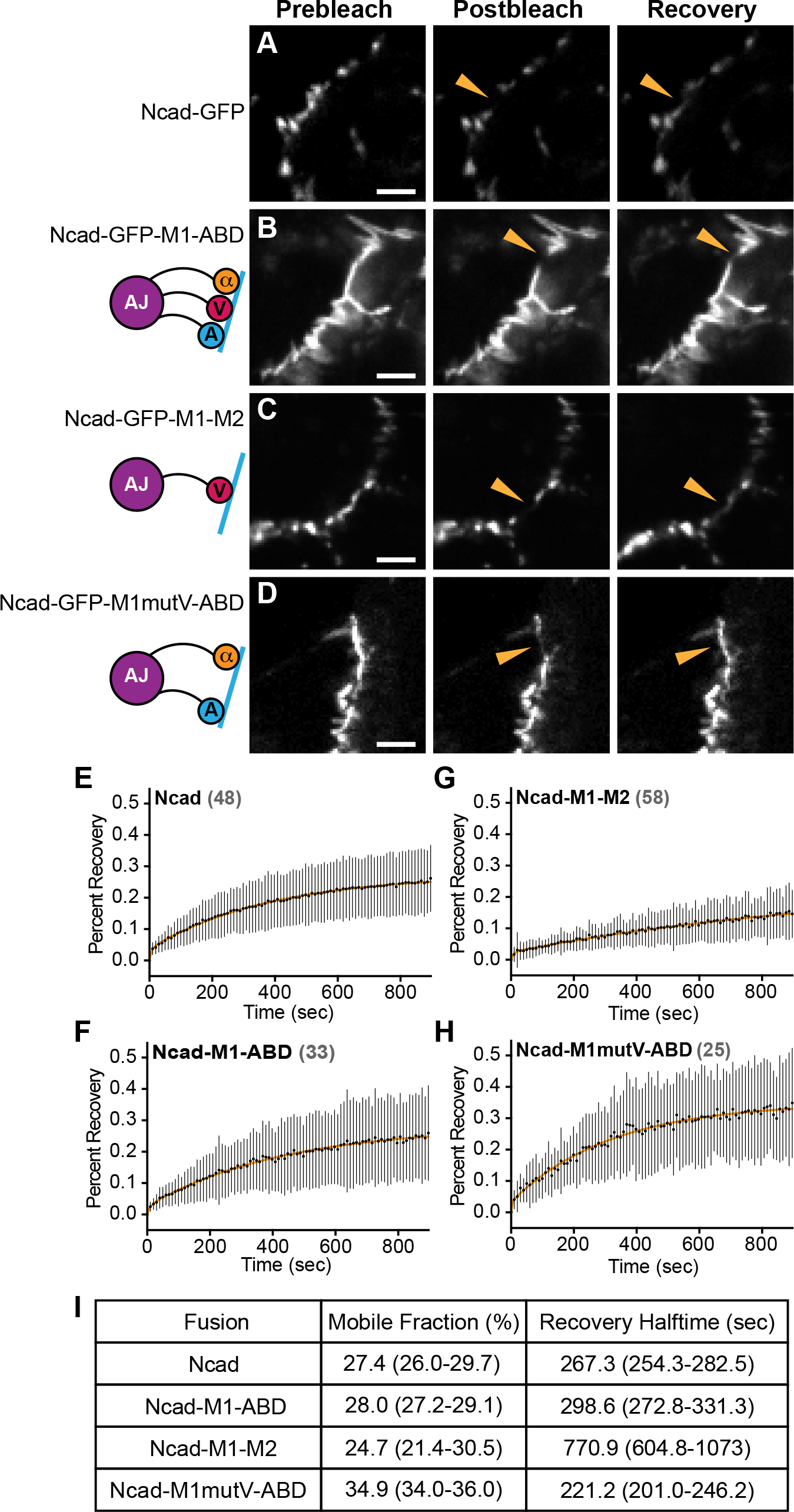
N-cadherin-GFP-αE-catenin fusion dynamics at cardiomyocyte cell-cell contacts. A-D. Representative prebleach, postbleach and recovery images from FRAP studies of Ncad^fx/fx^ cardiomyocytes infected with Cre and N-cadherin-GFP or N-cadherin-GFP-αE-catenin fusion adenoviruses. Orange arrowhead marks the FRAP region at a cell-cell contact. E-H. Plot of mean ± s.d. FRAP recovery fraction over 15 minutes. The mean is represented by a black circle and the standard deviation is shown as a black line. The data was fit to a double-exponential curve (orange line). The number of FRAP regions measured for each fusion construct is listed in grey. FRAP data was collected from at least 2 independent infections for each fusion. I. Summary of the mobile fraction (percentage) and recovery halftime (seconds). Scale bar is 5 μm in A-D.

We then analyzed the recovery rates of the mobile fraction slow pools. Ncad-GFP and Ncad-GFP-M1-ABD had similar recovery rates (Figure 5, B and G), consistent with the ability of Ncad-GFP-M1-ABD to reconstitute the AJ. Notably, Ncad-GFP-M1mutV-ABD had a recovery rate faster than Ncad-GFP, suggesting that vinculin regulates the dynamics of the AJ mobile pool (Figure 5, D and H). Consistent with this, Ncad-GFP-M1-M2, which binds vinculin constitutively, had a recovery rate that was nearly 3x slower than Ncad-GFP or Ncad-M1-ABD (Figure 5, C and F). We speculate that vinculin anchors the cadherin-catenin complex to actin to limit turnover of the mobile pool without affecting the immobile/mobile pool balance.

### Vinculin links the AJ to contractile myofibrils

We then questioned if the differences in cell-cell contact morphology and dynamics observed in cardiomyocytes expressing the Ncad-GFP-αEcat fusions could reflect fundamental changes in actin organization and/or linkage to the AJ. We used thin-section transmission electron microscopy to assess the ultrastructural organization of the AJ-actin interface. In adult mouse cardiac tissue, the ICD is a contorted, electron-dense structure where myofibrils are coupled between adjacent cells at AJs (Bennett *et al.*, 2006; Li *et al.*, 2019). In wild-type cultured cardiomyocytes, we observed a similar junctional morphology with electron-dense AJs joining myofibrils across cells (Figure 6, A and B, contact highlighted in purple). As expected, loss of N-cadherin dissolved cell junctions and prevented myofibril integration between neighboring cells (Figure 6, C and D). Importantly, AJ structure and myofibril pairing were rescued with N-cad-GFP expression in N-cadherin-null cells (Figure 6, E and F).

**Figure 6.**
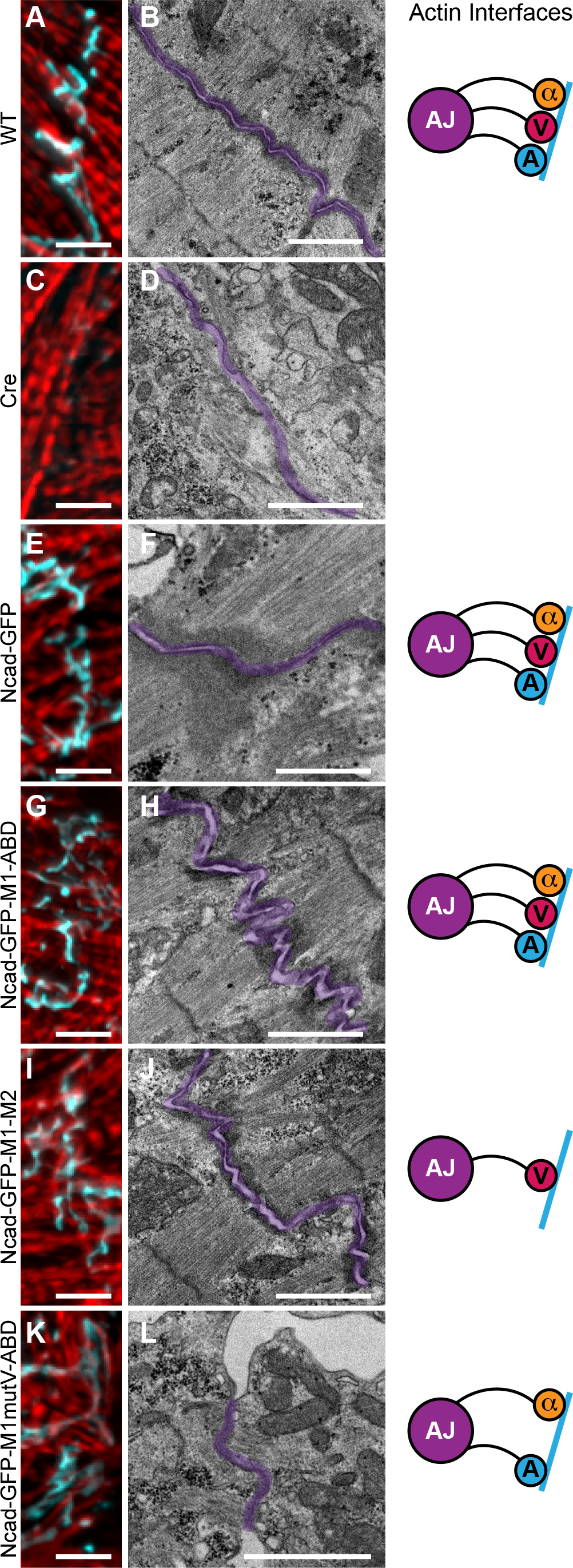
Vinculin recruitment is required to couple myofibrils to the AJ. Ncad^fx/fx^ cardiomyocytes uninfected (A, B), infected with Cre adenovirus (C, D), or infected with Cre and N-cadherin-GFP-αE-catenin fusion adenoviruses (E-L) were fixed and processed for staining (A, C, E, G, I, K) or thin section TEM (B, D, F, H, J, L). IF images are 2-3 μm deconvolved stacks of N-cadherin staining (A, C) or GFP signal from of N-cadherin-GFP-αE-catenin fusions (E, G, I, K) pseudo-colored blue and stained for F-actin (red). Representative TEM image shown from >60 images from at least three independent experiments. Cell-cell contacts are pseudo-colored purple. Scale bar is 5 μm in A-K; 1 μm in B, D, H, J, L; and 500 nm in F.

Next, we compared the ability of the Ncad-GFP-αE-cat fusions to organize actin along cardiomyocyte cell-cell contacts. Importantly, Ncad-GFP-M1-ABD restored myofibril coupling along thick, electron-dense junctions that were morphologically similar to controls (Figure 6, G and H; compare to Figure 6, A and B, E and F). As expected, lack of actin binding in Ncad-M1-M3 prevented cytoskeletal integration at cell-cell contacts (Supplemental Figure S3, A and B). Strikingly, Ncad-GFP-M1-M2, which connects N-cadherin to actin solely through vinculin, restored myofibril coupling and generated electron-dense junctions morphologically similar to Ncad-GFP-M1-ABD and controls (Figure 6, I and J). In marked contrast, constructs that could bind actin but were incapable of recruiting vinculin failed to restore normal contact morphology or myofibril coupling (Figure 6, K and L; Supplemental Figure S3, C-F). Ncad-GFP-M1mutV-ABD and Ncad-GFP-M2-ABD formed thin electron densities along elongated cell-cell contacts, but with little to no myofibril engagement. These results indicate that vinculin recruitment is required to link contractile actin to the cardiomyocyte AJ, as has been suggested in epithelia (Chen *et al.*, 2015). Neither the αE-catenin ABD nor afadin recruitment was sufficient to restore myofibril coupling. The increase in electron density along contacts with afadin recruitment suggests it may play a role in actin integration along contacts, but it is not sufficient to couple contractile actin in the absence of vinculin. Together, these results underscore the importance of vinculin in linking the AJ to contractile actin and highlight how αE-catenin coordinates cytoskeletal integration to provide mechanical connections between cells.

### Vinculin-binding ligands are not crucial to integration

Myofibrils are a highly specialized, contractile actin network, distinct from actin cables or stress fibers found in epithelial cells. However, the importance of vinculin in linking this unique network to the cardiomyocyte AJ is reminiscent of contractile actin linkages in epithelial cells. Vinculin anchors F-actin to the AJ (le Duc *et al.*, 2010; Yonemura *et al.*, 2010; Kale *et al.*, 2018) and can also recruit Ena/VASP proteins to promote actin assembly at junctions under tension (Leerberg *et al.*, 2014).To determine if cardiomyocytes use a similar linkage mechanism to epithelial cells, we probed for the Ena/VASP protein Mena (Krause *et al.*, 2003). Mena is recruited to epithelial contacts under tension (Leerberg *et al.*, 2014), is localized to the ICD in heart tissue (Aguilar *et al.*, 2011) and was identified in a proximity proteomics screen for N-cadherin-associated protein in cardiomyocytes (Li *et al.*, 2019). Immunostaining revealed limited recruitment of Mena to cell-cell contacts in cultured cardiomyocytes (Supplemental Figure S4A) that was lost after depletion of N-cadherin (Supplemental Figure S4B). Mena localization was only restored in fusion constructs that recruited vinculin (Supplemental Figure S4, D and E vs. S4F). However, we observed no difference in Mena recruitment between Ncad-M1-M2 and Ncad-M1-ABD despite Ncad-M1-M2 enriching vinculin at cell-cell contacts (Figure 4L). Thus, while Mena may function in linking contractile F-actin to cardiomyocyte AJs, its recruitment is limited and not correlated with vinculin levels, suggesting a more peripheral role in regulating cardiomyocyte junctional actin. α-Actinin binds vinculin (Kroemker *et al.*, 1994) and αE-catenin (Knudsen *et al.*, 1995; Nieset *et al.*, 1997). α-Actinin crosslinks actin filaments at myofibril Z-discs and is critical for cardiomyocyte organization (Frank and Frey, 2011). We did not observe any changes in α-actinin localization with a loss of N-cadherin (Supplemental Figure S5A-B). However, in Ncad-M1-M2, we observed a modest recruitment of α-actinin to contacts (Supplemental Figure S5D). We were not able to determine if α-actinin was recruited through vinculin and/or αE-catenin M1-M2. While increased α-actinin recruitment could impact Ncad-M1-M2 dynamics (Figure 5C, F), enrichment is not required for AJ coupling to F-actin.

### Ligand requirements differ for junctional complex assembly

We then wanted to determine if the Ncad-GFP-αEcat fusions could restore the two other major junctional complexes at the ICD: gap junctions and desmosomes. Gap junctions electrically couple cardiomyocytes and their formation is predicated on cadherin localization to cell-cell contacts (Noorman *et al.*, 2009). N-cadherin depletion causes loss of Cx43, the pore-forming protein of gap junctions (Figure 2K), and Cx43 localization to cell-cell junctions can be restored by expressing Ncad-GFP (Figure 7A). Importantly, Cx43 contact localization was restored with all Ncad-GFP-αEcat fusions (Figure 5B-D; Supplemental Figure S6A-C) independent of ligand recruitment or actin engagement. This indicates that N-cadherin delivery to the plasma membrane is sufficient to localize Cx43 to nascent contacts, though gap junction stabilization may require ligand recruitment (Zemljic-Harpf *et al.*, 2014).

**Figure 7.**
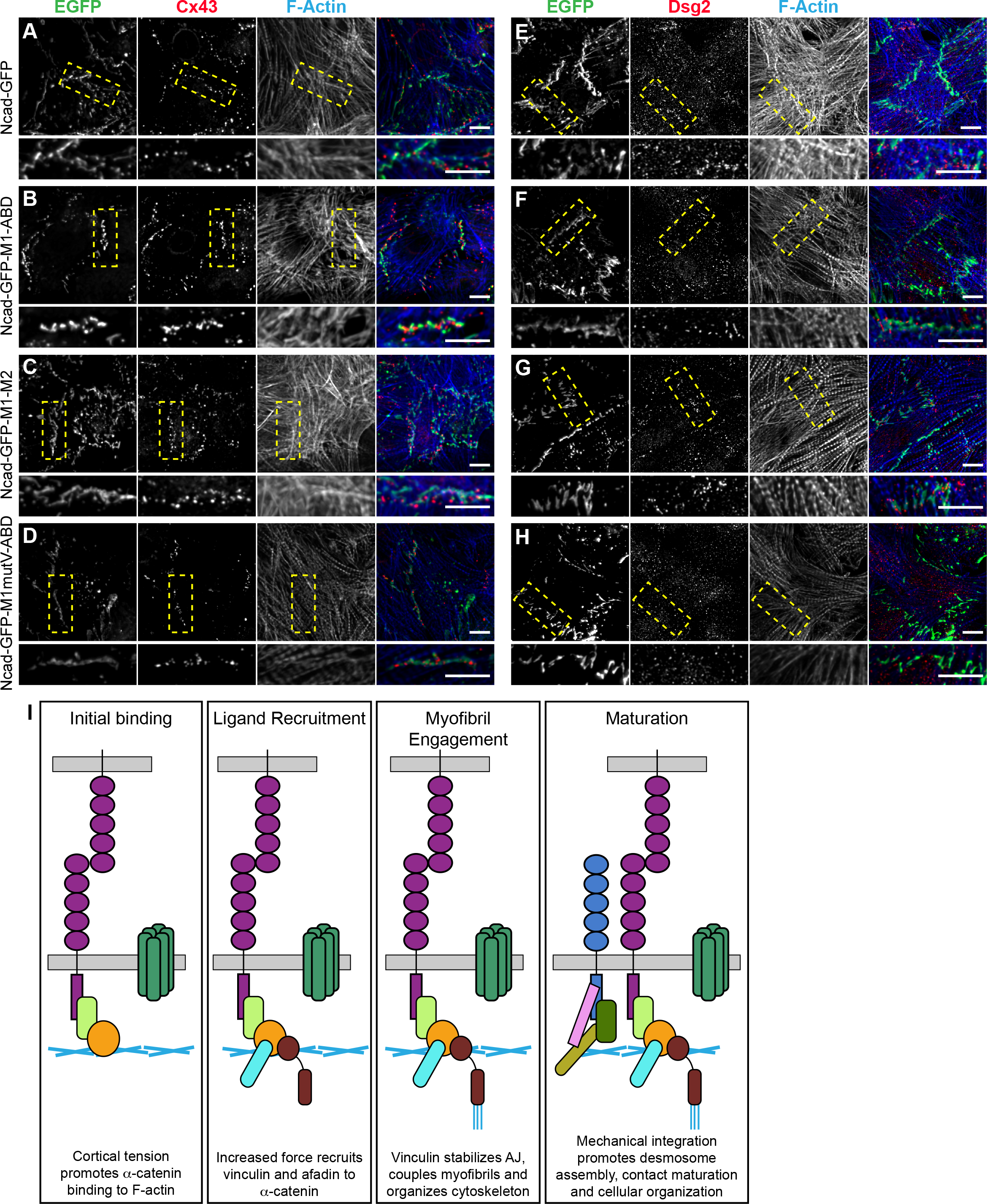
N-cadherin-GFP-αE-catenin fusions restore junctional complexes. N-cadherin-null cardiomyocytes infected with N-cadherin-GFP (A, E) or N-cadherin-GFP-αE-catenin fusion adenoviruses (B-D, F-H). A-D. Cells were fixed and stained for connexin 43 and F-actin. Individual and merged GFP (green), connexin 43 (Cx43, red) and F-actin (blue) channels shown. E-H. Cells were stained for desmoglein 2 and F-actin. Individual and merged GFP (green), desmoglein 2 (Dgs2, red) and F-actin (blue) channels shown. Images are a max projection of 2-3 μm deconvolved stacks. Bottom image is a magnification of boxed contact. Scale bar is 10 μm. I. Schematic of AJ integration with cardiomyocyte actin networks and contact maturation.

Desmosomes also require AJ establishment for assembling along cell-cell contacts (Nekrasova and Green, 2013). Recent works suggests that E-cadherin recruits desmoglein 2 (Dsg2) through direct extracellular *cis* interactions to promote desmosome assembly at nascent contacts in epithelial cells (Shafraz *et al.*, 2018). We assessed the ability of the fusion constructs to restore desmosome recruitment after loss of N-cadherin. Interestingly, Dsg2 recruitment was only observed in fusions that could recruit vinculin (Figure 7E-G vs. H; Supplemental Figure S6D-F). These data imply that vinculin recruitment could provide a needed mechanical linkage to drive desmosome assembly at nascent cardiomyocyte contacts.

## Discussion

Together, our results provide novel insights into the AJ-myofibril linkage in cardiomyocytes. We show that vinculin recruitment through α-catenin is required to couple the cardiomyocyte AJ to contractile actin and promote AJ stability and contact maturation. The mechanical properties of α-catenin thus function to integrate cytoskeletal networks at cardiomyocyte cell-cell contacts to build force-resilient junctions.

Vinculin is recruited to epithelial AJs in tension-dependent manner where is thought to help anchor the AJ to actin (le Duc *et al.*, 2010; Yonemura *et al.*, 2010). In addition to creating a new linkage between F-actin and the AJ, vinculin can also recruit ligands such as Mena to promote actin assembly at junctions under tension (Leerberg *et al.*, 2014). While Mena localizes to cardiomyocyte cell-cell contacts, we did not observe a concomitant enrichment of Mena with constitutive vinculin recruitment to cardiomyocyte AJs (Fig. S5). We did not detect VASP at cardiomyocyte AJs (data not shown). Thus, our results suggest that vinculin-mediated ligand recruitment may not be the primary driver of increased stability and myofibril integration at the AJ. Instead, the actin binding domain of vinculin may play a critical role in coupling the AJ to contractile actin.

Myofibrils arrange their actin filaments so that the myosin motors exert force as they move toward the barbed end. Recent biophysical data has shown that the vinculin-actin interaction is asymmetrical, where the bond is strengthened when an actin filament is under pointed (-) end-directed load (Huang *et al.*, 2017). Additional work has demonstrated that differential recruitment of vinculin to sites of high tension in epithelial cells is used to balance tensile and shear forces across cell contacts (Kale *et al.*, 2018). We propose a model in which tension activates αE-catenin at nascent cardiomyocyte contacts to promote ligand binding (Fig. 7I). Vinculin recruitment, in turn, promotes myofibril binding to strengthen the AJ and orchestrate junctional maturation and mechanical integration. We speculate that vinculin recruitment creates a self-amplifying tension feedback loop to promote junctional planar organization necessary for heart muscle function. Thus, vinculin functions as both a mechanical linchpin and critical organizer of actomyosin and AJ architecture to regulate cardiomyocyte adhesion. We suggest that this is a general mechanism cells use to organize contractile actin networks across various cells types. For example, vinculin is enriched at tricellular junctions in epithelial sheets where contractile actin terminates perpendicularly to AJs (Higashi and Miller, 2017), similar to cardiomyocytes. Linking to specific actin networks through selective ligand recruitment would allow the AJ to control mechanical load and respond to changes in cellular tension.

Vinculin is a mechano-responsive protein found at both cell-cell and cell-ECM contacts (Bays and DeMali, 2017). Failing human cardiac tissue shows an increased expression of vinculin though with a less organized localization pattern (Heling *et al.*, 2000). Aging non-human primate and mouse models show increased vinculin expression compared to younger animals, with increased localization at both the ICD and cell-ECM adhesions (Kaushik *et al.*, 2015). Aging, and ultimately failing, cardiac tissue shows signs of fibrosis resulting in an increased ECM stiffness (Biernacka and Frangogiannis, 2011). Increased ECM stiffness has been linked to molecular remodeling of myofibril integration at cell-cell versus cell-ECM adhesions, causing a decrease of force propagation across adhered cardiomyocytes (McCain *et al.*, 2012). Interestingly, we found that M1-M2 expression caused a loss of vinculin at cell-ECM contacts (data not shown). Consistent with this, vinculin can be selectively enriched at cell-cell contacts or cell-ECM contacts when external tension is applied to either of these areas individually (le Duc *et al.*, 2010; Plotnikov *et al.*, 2012). Our results highlight the critical role of vinculin at cardiomyocyte AJs and provide a possible explanation for how changes in ECM stiffness and concomitant cell-ECM adhesion expansion would disrupt the balance of vinculin to impair myofibril integration and decrease cardiac function.

## Materials and Methods

### Plasmids

To build the N-cadherin-GFP-αE-catenin fusions, αE-catenin fragments encoding aa273-510, aa273-651, and aa273-906 were cloned into pEGFP-C1 by PCR. Next, Gibson Assembly (NEBuilder HiFi DNA Assembly Kit, New England Biolabs) was used to clone the N-cadherin fragment aa1-839 into pcDNA3.1 (Thermofisher). Gibson assembly was then used to insert the EGFP-αE-catenin fragments into the pcDNA3.1 N-cadherin aa1-839 backbone, downstream and in-frame with N-cadherin. During construction, a 12 aa glycine and alanine linker was inserted between N-cadherin and αE-catenin to increase flexibility.

The point mutations R329A, R330A, L347A, L348A and Y351A were introduced by site directed mutagenesis (Agilent) in the αE-catenin M-region to inhibit vinculin binding (Chen *et al.*, 2015).

### Cardiomyocyte Isolation and Culture

All animal work was approved by the University of Pittsburgh Division of Laboratory Animal Resources. Outbred Swiss Webster mice were used to generate wild-type cardiomyocytes for blebbistatin experiments. N-cad^fx’fx^ conditional knockout mice (Jackson Labs, stock #007611, (Kostetskii *et al.*, 2005) were used to generate N-cadherin null cardiomyocytes.

Tissue culture dishes or MatTek dishes (35 mm dish with 10 mm microwell) were coated with rat tail Type I collagen (Millipore) diluted to 0.5 μg/μl in PBS for 30 minutes at room temperature. Dishes were dried and treated with UV radiation for 1 hour, after which they were washed with PBS, dried and stored at room temperature in the dark.

Neonatal mouse cardiomyocytes were isolated as described (Ehler *et al.*, 2013). Briefly, mouse pups were sacrificed at P1-P3, the hearts were removed, cleaned, minced, and digested overnight at 4°C in 20 mM BDM (2,3-Butanedione monoxime) and 0.0125% trypsin in HBSS. The following day, heart tissue was digested further in15 mg/mL Collagenase/Dispase (Roche) in Leibovitz media with 20 mM BDM to create a single cell suspension. Cells were pre-plated for 1.5-2 hours in plating media (65% high glucose DMEM, 19% M-199, 10% horse serum, 5% FBS and 1% penicillin-streptomycin) to remove fibroblasts and endothelial cells Cardiomyocytes were plated on MatTek dishes (1.5×10^5^) or 12-well dishes (4.5×10^5^) in plating media. 16 hours post-plating, the plating media was exchanged for maintenance media (78% high glucose DMEM, 17% M-199, 4% horse serum, 1% penicillin-streptomyocin, 1 μM AraC, and 1 μM Isoproternol).

### Adenovirus Production and Infection

N-cadherin-GFP-αE-catenin fusions were expressed as adenoviruses using the AdEasy System as described (Luo *et al.*, 2007; Li *et al.*, 2019). Briefly, N-cadherin-EGFP-αE-catenin fusions were moved in to pShuttle-CMV using NEBBuilder HiFi DNA Assembly Master Mix (New England Biolabs). Positive clones were transformed into AdEasier *E. coli* cells to generate recombinant adenovirus DNA. Adenoviral plasmids were then transfected into HEK293 cells for virus production. Virus was amplified and purified using AdenoPACK 20 Adenovirus (Ad5) purification & concentration kit (Sartorius).

Virus titer was determined by quantitative Polymerase Chain Reaction (qPCR) using Adeno-X qPCR Titration Kit (Clontech) on an Applied Biosystems 7900HT.

Adenovirus expressing Cre (Ad(RGD)-CMV-iCre) was purchased from Vector Biolabs.

Cdh2^fx/fx^ cardiomyocytes were infected with adenovirus Cre at MOI (multiplicity of infection) 75 on the day of plating to achieve 100% infection. 16 hours after virus addition, the media was replaced with maintenance media. 6-8 hours later (22-24 hours after Cre infection) cardiomyocytes were infected with the N-cadherin-GFP-αE-catenin fusion adenovirus at MOI 10-15 to achieve >50% infection rate. Cardiomyocytes were fixed 96 hours after Cre infection for analysis.

### Immunofluorescence

Cells were processed for immunofluorescence as follows: cells were fixed in warmed (37°C) 4% EM grade paraformaldehyde in PHM buffer (60 mM PIPES pH 7.0, 25 mM HEPES pH 7.0, 2 mM MgCl_2_ and 0.12 M Sucrose) for 10 minutes and washed twice with PBS. Cells were permeabilized with 0.2% Triton X-100 in PBS for 4 minutes and washed twice with PBS. Cells were blocked in 10% BSA (Sigma) in PBS for 1 hour at room temperature. Samples were incubated with primary antibodies in PBS + 1% BSA for 1 hour at room temperature, washed 2X in PBS, incubated with secondary antibodies in PBS + 1% for 1 hour at room temperature, washed 2X in PBS and then mounted in Prolong Diamond (Thermo Fisher Scientific). All samples were cured at least 24 hours before imaging.

For blebbistatin experiments, cardiomyocytes (96 hours post-plating) were treated with 100 μM blebbistatin in DMSO or DMSO control or 10 minutes to 1 hour. Cells were incubated at 37°C during treatment. After incubation, cells were first pre-permeabilized in 0.2% Triton X-100 in PBS for 2 minutes, then fixed and labeled as described.

### Antibodies

Primary antibodies used for immunostaining were: anti-αE-catenin (1:100; Enzo Life Sciences ALX-804-101-C100), anti-β-Catenin (1:250; BD Transduction Laboratories 610154), anti-Plakoglobin (1:100; Cell Signaling 2309), anti-Vinculin (1:800; Sigma Aldrich V9131), anti-N-cadherin (1:250; Invitrogen 99-3900), anti-l-Afadin (1:500; Sigma Aldrich A0349), anti-Connexin-43 (1:100; ProteinTech 15386-1-AP), anti-Plakophilin 2 (1:10; Progen 651101), anti-Desmoglein 2 (1:250, Abcam EPR6768), anti-α-Actinin (1:250, Sigma A7811), anti-Mena (1:300, mouse monoclonal, a kind gift from Frank Gertler) and anti-Cre Recombinase (1:300, Cell Signaling 12830). Secondary antibodies used were goat anti-mouse or anti-rabbit IgG conjugated to Alexa Fluor-488, 568, or 647 (1:250; Invitrogen). F-actin was visualized using an Alexa Fluor dye conjugated to phalloidin (1:100, ThermoFisher Scientific).

### Whole Cell Lysis and Immunoblotting

Cardiomyocytes were cultured on collagen-coated 12-well dishes (see above). 96 hours after plating, cardiomyocytes were lysed with RIPA buffer supplemented with 1X protease inhibitors (Millipore). Lysate protein concentration was determined by BCA Assay (BioRad). 15 μg of lysate was loaded per well and resolved on a 10% SDS PAGE and then transferred to a PVDF membrane. The membrane was blocked in 5% BSA in 1X TBST with 0.02% NaN_3_ for 1 hour at room temperature. Primary antibodies (N-cadherin 1:2500, GAPDH 1:750 (Abcam ab9485)) were diluted in 5% BSA in 1X TBST with 0.02% NaN3 overnight at 4°C, followed by three 15 minute TBST washes. LI-COR secondary antibodies (Goat anti-mouse 680; goat anti-rabbit 800, 1:15,000) were diluted in 1X TBST with 0.02% NaN3 and incubated for 1 hour at room temperature. The membrane was washed three times in 1X TBST and once in PBS. The membrane was imaged on a LI-COR Odyssey imaging system. Band intensities were quantified in ImageJ and plotted in Prism (GraphPad).

### Confocal Microscopy

Cells were imaged with a 100X 1.49 NA objective or a 40X 1.30 objective on a Nikon Eclipse Ti inverted microscope outfitted with a Prairie swept field confocal system, Agilent monolithic laser launch and Andor iXon3 camera using NIS-Elements (Nikon) imaging software. Maximum projections of 2-3 um image stacks were created and deconvolved (3D Deconvolution) in NIS-Elements (Nikon) for presentation. Expression and staining levels were adjusted for presentation purposes in Photoshop (Adobe). All levels were corrected the same across each figure except Figure 6 where the phalloidin labeling of F-actin was modified individually to account for differences in staining and in Figure 5 to account for changes in focal plane/expression for live cell imaging. Note that Ncad-GFP-ABD levels were adjusted individually in Supplemental Figures S2, S3, and S6 as this construct localized to cell-cell contacts less efficiently than the other fusions.

### FRAP experiments

FRAP experiments were conducted on a Nikon swept field confocal microscope (describe above) outfitted with a Tokai Hit cell incubator and Bruker miniscanner. Actively contracting cells were maintained at 37°C in a humidified, 5% CO_2_ environment. User-defined regions along cell-cell contacts were bleached with a 488 laser and recovery images collected every 10 seconds for 15 minutes. FRAP data was quantified in ImageJ (NIH) and average recovery plots were measured in Excel (Microsoft). All recovery plots represent data from two independent transfections of unique cell preps. The data were fit to a double-exponential curve to determine the mobile fraction and half time of recovery in Prism (Graphpad). Only recovery rates of the slow pool are reported as this was the dominant mobile pool (87-91%) for all constructs.

### Electron Microscopy

Cardiomyocytes were grown on collagen-coated MatTek dishes and fixed as described above. After fixation and washing, cells were incubated with 1% OsO4 for one hour. After several PBS washes, dishes were dehydrated through a graded series of 30% to 100% ethanol, and then infiltrated for 1 hour in Polybed 812 epoxy resin (Polysciences, Warrington, PA). After several changes of 100% resin over 24 hours, cells were embedded in inverted Beem capsules, cured at 37°C overnight and then hardened for 2 days at 65°C. Blocks were removed from the glass dish via freeze/thaw method by alternating liquid Nitrogen and 100°C water. Ultrathin (60nm) sections were collected on to 200-mesh copper grids, stained with 2% uranyl acetate in 50% methanol for 10 minutes and 1% lead citrate for 7 minutes. Samples were photographed with a JEOL JEM 1400 PLUS transmission electron microscope (Peabody, MA) at 80kV with a Hamamatsu ORCA-HR bottom mount camera.

### Image Analysis

Vinculin and afadin recruitment to the N-cadherin-EGFP-αE-catenin fusion constructs was analyzed in ImageJ. A single plane was selected from the z-stack where the contact was most in focus and IsoJ Dark thresholding was used to create a mask of the EGFP channel to define the region of analysis (cell-cell contacts). The vinculin or afadin signal intensity was then measured within the masked region. Next, three random intensity measurements of vinculin or afadin were taken in the cell cytoplasm and these values were averaged. Finally, the ratio of vinculin or afadin intensity within the mask was divided by the cytoplasmic signal to calculate the contact/cytoplasmic ratio. Colocalization data were plotted with Prism software (GraphPad). A One-way ANOVA with multiple comparisons was performed; p<0.05.

Blebbistatin experiments were analyzed in a similar method. A single plane was selected from the z-stack where the contact was most in focus. IsoJ Dark thresholding was used to create a mask of the tension-insensitive marker, and this mask was applied to either the vinculin or afadin channels to determine their intensity at the contact. Intensity values of treated (blebbistatin) and control (DMSO) samples were plotted in Prism software (GraphPad).

## Acknowledgements

We thank members of the Kwiatkowski lab for their helpful comments on the manuscript.

